# Reproductive developmental dynamics during the early and intermediate prepuberal periods in Nelore breed (*Bos indicus*) calves

**DOI:** 10.1101/2021.12.02.470948

**Authors:** Taynan Stonoga Kawamoto, João Henrique Moreira Viana, Maurício Machaim Franco, Otávio Augusto Costa de Faria, Andrei Antonioni Guedes Fidelis, Ricardo Alamino Figueiredo

**Affiliations:** Department of Veterinary, Federal University of Uberlandia, Uberlandia, MG, Brazil; Animal Reproduction Laboratory, Embrapa, Genetic Resources and Biotechnology, Brasília, DF, Brazil; Department of Veterinary, University of Brasilia, Brasília, DF, Brazil

**Author notes:** Animal Reproduction Laboratory, Embrapa, Genetic Resources and Biotechnology, Brasília, DF, Brazil. Corresponding author: (JHMV).

**Keywords:** *Bos indicus*, puberty, reproductive physiology, rump development

## Abstract

The demand for calves as oocyte donors for in vitro embryo production is growing. However, Bos indicus have a late puberty, and some aspects of the reproductive physiology during the prepubertal period remain unclear. We characterized endocrine and morpho-functional reproductive features in Nelore calves (n=8) at 2-5 (early prepubertal period, EPP) and from 8-11 months old (mo., intermediate prepubertal period, IPP). The calves’ ovaries and uterus were B-mode transrectal ultrasonography examined, and blood samples were collected every second week. The antral follicles number and size, and ovarian and uterine horn diameters, were recorded, and plasma FSH and LH concentrations were measured (RIA). Non-pregnant, non-lactating cyclic Nelore cows (n=8) were used as controls for endocrine endpoints. Somatic development was monitored by monthly weighing, and 3D scanning of the rump area. The somatic and endocrine endpoints were compared within and between EPP and IPP, and between each period and control cows. Associations were determined by the Spearman correlation method, and the developmental rates were determined by non-linear regression. All morphological endpoints, except antral follicle count, increased (P < 0.001) from the EPP to the IPP. However, within each period differences occurred only at EPP. During the EPP LH and FSH plasma concentrations were similar (P > 0.05), whereas during the IPP LH was lower (P < 0.05) and FSH was higher (P < 0.001) than control cows. The EPP calves showed moderate to high positive correlations among ovarian, uterine, and somatic endpoints. Conversely, the IPP such correlations were mostly weak. In summary, distinct ovarian activity and development patterns of primary and secondary sexual characteristics occurred in Nelore calves at 2-5 mo compared to 8-11 mo.

## Introduction

The period from birth until puberty is marked by a number of changes that will lead to the maturity of the hypothalamus-hypophysis-ovary axis and prepare the reproductive system for the successful generation of viable offspring. In cattle, as in many other species, the ovary is active during this period, and follicle recruitment and growth up to the antral phase will occur [1–4]. Sexual steroids produced by growing ovarian follicles promote the development of the tubular part of the reproductive system, but they also play a crucial role in somatic development, acting in fat deposition [5] and in muscle [reviewed by 6] and bone development [5, 7, 8]. However, final follicle maturation and subsequent ovulation will not occur before puberty, mainly due to a higher sensitivity of the hypothalamus to the negative feedback of ovarian estradiol [9], which prevents further increases in FSH and LH.

In the past, studies on cattle puberty focused on the physiology of the events during the peripubertal period, particularly from 8 mo on, aiming to reduce the age at puberty [10]. However, the possibility of generating offspring from prepubertal cattle by the use of *in vitro* embryo production (IVEP) technologies renewed the interest in understanding the reproductive physiology during earlier prepubertal phases. The production of offspring from prepubertal cattle by IVEP has been reported since the 1990s [11, 12]. However, low embryo rates and the technical difficulties for oocyte recovery overcame the economic advantages of using donors with genetic merit still undetermined. This scenario changed with the recent advances in genomics and the development of commercial microchips to evaluate the genomic-predicted transmitting ability (GPTA) for type and production traits in different cattle breeds [13]. Selection of genetically superior cattle can now be determined from birth [13], and this has boosted the demand for the use of prepubertal calves as oocyte donors.

Two main techniques are currently used to recover oocytes from prepubertal female cattle, namely laparoscopic and ultrasound-guided follicle aspiration (LOPU and OPU, respectively). The LOPU is used mainly in calves up to 5 mo, when rump development limits the rectal access to the ovaries. On the other hand, OPU has been the technique of choice for older calves and prepubertal heifers [14]. In this regard, the sequence of events that occur in these early time windows are key to understand how to improve the developmental potential of oocytes recovered from calves. Despite the occurrence of antral follicle growth before puberty, oocytes recovered from prepubertal calves have lower developmental potential compared to those recovered from pubertal heifers or cows [15, 16]. The causes underlying this difference are not fully understood, but changes in gonadotropin stimulation and in subsequent steroid production play a central hole in oocyte developmental potential. Transplantation of grafts from calf ovary to adult cows restores oocyte potential [9], whereas treatment with exogenous FSH or eCG increases the blastocyst rates obtained with calf oocytes [10].

The prepubertal period can be subdivided in phases characterized by distinct morphological and endocrine features. In male calves, the infancy period lasts up to 3 mo and is associated with a low LH secretion. After that, LH pulse frequency increases throughout the prepubertal period. In females, ovary size, antral follicle count (AFC), and dominant follicle (DF) maximum size increase up to 4 mo, plateau between 4 and 8 mo, and resume increase until puberty [1]. It should be noted that, however, most studies addressing the reproductive physiology during the prepubertal period were done in *Bos taurus* breeds [1, 16, 17]; *B. taurus indicus* cattle is less precocious than European breeds (*B. Taurus* [18, 19]), with significant differences in ovarian physiology, such as a higher antral follicle count (AFC) and a lower diameter of the dominant follicles [20]. Thus, it is likely that the timing or the magnitude of the events that will take place in the early prepubertal period may also differ from those reported for *B. taurus*.

The aim of the present study was to investigate the endocrine and morpho-functional reproductive characteristics of Nelore calves in two time windows, namely at the early and intermediate prepubertal periods. We hypothesized that these periods are characterized by distinct patterns of ovarian activity, which in turn drive reproductive and somatic development patterns in each phase.

## Materials and Methods

### Animals and location

This study was conducted at the Sucupira Experimental Farm, Embrapa Genetic Resources and Biotechnology, Brasilia, Brazil. The contemporaneous Nelore calves (n = 8) used in the present study were generated by timed artificial insemination (TAI) with X-sorted semen. For endocrine endpoints, non-pregnant and non-lactating pluriparous cows (n = 8) from the same herd were used as controls. Calves were raised under pasture (*Brachiaria decumbens*), with *ad libitum* access to water and minerals. Weaning occurred at 7 mo. The average daily weight gain during the experiment was 0.57 ± 0.2 kg/day (from an average of 90.9 ± 8.3 kg BW at the beginning to 245.0 ± 12.3 kg BW at the end of the experiment).

### Experimental design

Two evaluation periods were defined: 2 to 5 mo, referred to as early prepuberal period (EPP), and 8 to 11 mo, referred to as intermediate prepubertal period (IPP). These time windows were chosen to represent the initial and intermediate prepubertal periods, taking in account the expected age at puberty in the Nelore breed (22 to 36 mo, [21]) as well as the ages at which LOPU and OPU are usually performed in calves. In each period, calves were examined weekly by transrectal ultrasonography to measure ovarian and uterine horn diameters (mm), total antral follicle count (AFC), the number of follicles ≤ or > 4 mm, and the diameter of the largest follicle (mm). Blood samples were collected once a day for FSH and twice a day for LH plasma evaluations, at the same day of the ultrasonographic exams. Once a month, the calves were weighted and underwent rump biometric measurements. When calves reached 15 mo, monthly ultrasound scanning was used to determine the age at puberty, based on the first detection of a corpus luteum.

### Ultrasonographic evaluations

Ovarian and uterine evaluations were performed using a B-mode ultrasound (MyLab 30 VetGold, Esaote. Genova, Italy) equipped with a 5- to 7.5-MHz linear rectal probe. For all exams, settings related with frequency (7.5 MHz), focus depth, and gain adjustments were standardized. The AFC and the number of follicles ≤ or > 4 mm were determined visually during ovarian scanning. Linear measurements of the diameter of the ovary, of the largest follicle present, and of the uterine horns were done using the internal caliper of the ultrasound. During the first evaluation period (2 to 5 mo, EPP), a probe guide was used to enable transrectal examination without rectal manipulation of the genital tract, whereas for the second period (8 to 11 mo, IPP), as well as for cows, conventional transrectal scanning procedures were used.

### Blood samples and hormonal measurements

Blood samples were collected by jugular vein puncture using 21G, double-lumen needles and vacuum tubes with EDTA. The samples were kept on ice until being centrifuged (3,000 x g for 15 min), and 1-mL aliquots of plasma were stored at −20°C. The LH and FSH analyses were performed by radioimmunoassay (RIA) as described by [22] at the Laboratory of Endocrinology of the Sao Paulo State University (UNESP, Araçatuba, SP, Brazil). Assay sensitivity was 0.126 ng/mL for LH and 0.390 ng/mL for FSH. The intra-assay coefficient of variation was < 11%.

### Rump biometric data

Rump width, length, and geometry were calculated using as reference points the prominences of the *tuber ischiae* and the *tuber coxae* of the pelvic bones. Three-dimensional images of the rump area were acquired by structured infrared light scanning, using a portable sensor (iSense, 3D Systems, Rock Hill, SC, USA) connected to a tablet computer (iPad Air 2, Apple Inc., Cupertino, CA, USA). This computer was equipped with a real-time scanning app (https://itunes.apple.com/us/app/isense/id807510940). The nominal resolution of this equipment at 0.5 m distance was 0.9 mm for the x/y axes and 1.0 mm for the z axis (depth). The points cloud data were transformed in a geometric surface and stored as OBJ files. The 3D images were then edited using the open-source software MeshLab (SourceForge, USA) to delete unspecific scans from the nearby objects and to perform the measurements. As a validation procedure, manual measurements were taken using a metric tape. There were no significant differences in rump width or length, as measured directly with scales or obtained in the 3D images, and only the latter were used in this study. The rump area was defined as the area of the trapezium formed by the width at the ilium, the width at the ischium, and the rump length. Changes in rump geometry were demonstrated in snapshots taken from the back view of 3D images from calves at 2, 5, 8, and 11 mo. The trapeziums were drawn using the same reference points as for biometric measurements.

### Statistical analysis

Data of continuous outcome variables were first examined for normality and homogeneity of variances using the Shapiro-Wilk and Bartlett tests. Variables with normal distribution were evaluated by ANOVA, differences between parity groups (calves vs. cows) or between prepubertal moments (EPP vs. IPP) were compared using the Student t test, otherwise analyzed using the Mann-Whitney test. The association between reproductive and biometric data was evaluated using Spearman’s correlation. The reproductive developmental rate was evaluated by non-linear regression analysis (Gompertz curve), and the validation of this curve was tested by the deviance of the fitted curve compared to a straight line. The Gompertz model equation was Z *exp (-b*exp (a*x), where “Z” is the maximum data value, “b” is the integration constant, and “a” corresponds to the acceleration rate. This non-linear regression analysis (Gompertz curve) was also used to compare rump geometry changes at different ages. All statistical procedures were performed using the GraphPad Prism (v. 6) or the R (v. 3.6.1) software, and a P value ≤ 0.05 was considered as statistically significant. Results are shown as mean ± SEM.

## Results

The results of the ultrasonographic evaluation of the calves’ ovaries and uterus during the EPP and IPP are shown in Table 1. The ovarian and the uterine diameter, the number of follicles larger than 4 mm, and the diameter of the largest follicle were greater in IPP calves compared to EPP calves (P < 0.001). However, there was no difference (P > 0.05) in AFC between EPP and IPP. The EPP calves had LH and FSH plasma concentrations similar (P > 0.05) to cows, whereas IPP calves had lower LH (P < 0.05) and higher FSH (P < 0.001) concentrations than cows (Fig 1, A-D).

**Table 1.** Average values (mean ± SD) of reproductive and somatic endpoints in Nelore calves during two time windows in the prepubertal period: from 2 to 5 and from 8 to 11 months old.

| Endpoint | Age group |  | P value |
| --- | --- | --- | --- |
|  | 2–5 months | 8–11 months |  |
| Ovarian diameter (mm) | 16.5 ± 2.4 <sup>A</sup> | 18.1 ± 2.1 <sup>B</sup> | < 0.0001 |
| Antral follicle count (n) | 20.7 ± 6.7 | 22.0 ± 8.9 | 0.5412 |
| Number of follicles > 4mm | 3.4 ± 1.6 <sup>A</sup> | 4.9 ± 2.2 <sup>B</sup> | < 0.0001 |
| Diameter of the largest follicle (mm) | 7.2 ± 1.5 <sup>A</sup> | 7.9 ± 1.3 <sup>B</sup> | 0.0001 |
| Uterine diameter (mm) | 8.4 ± 1.0 <sup>A</sup> | 10.1 ± 1.0 <sup>B</sup> | < 0.0001 |
| Average body weight | 118.8 ± 22.9 <sup>A</sup> | 227.0 ± 18.7 <sup>B</sup> | < 0.0001 |
| Mean rump area (cm<sup>2</sup>) | 514.2 ± 101.4 | 856.7 ± 79.1 | ----- |

Different letters in the same line are different. T-test and Mann-Whitney analysis, P<0.05.

**Fig 1.**
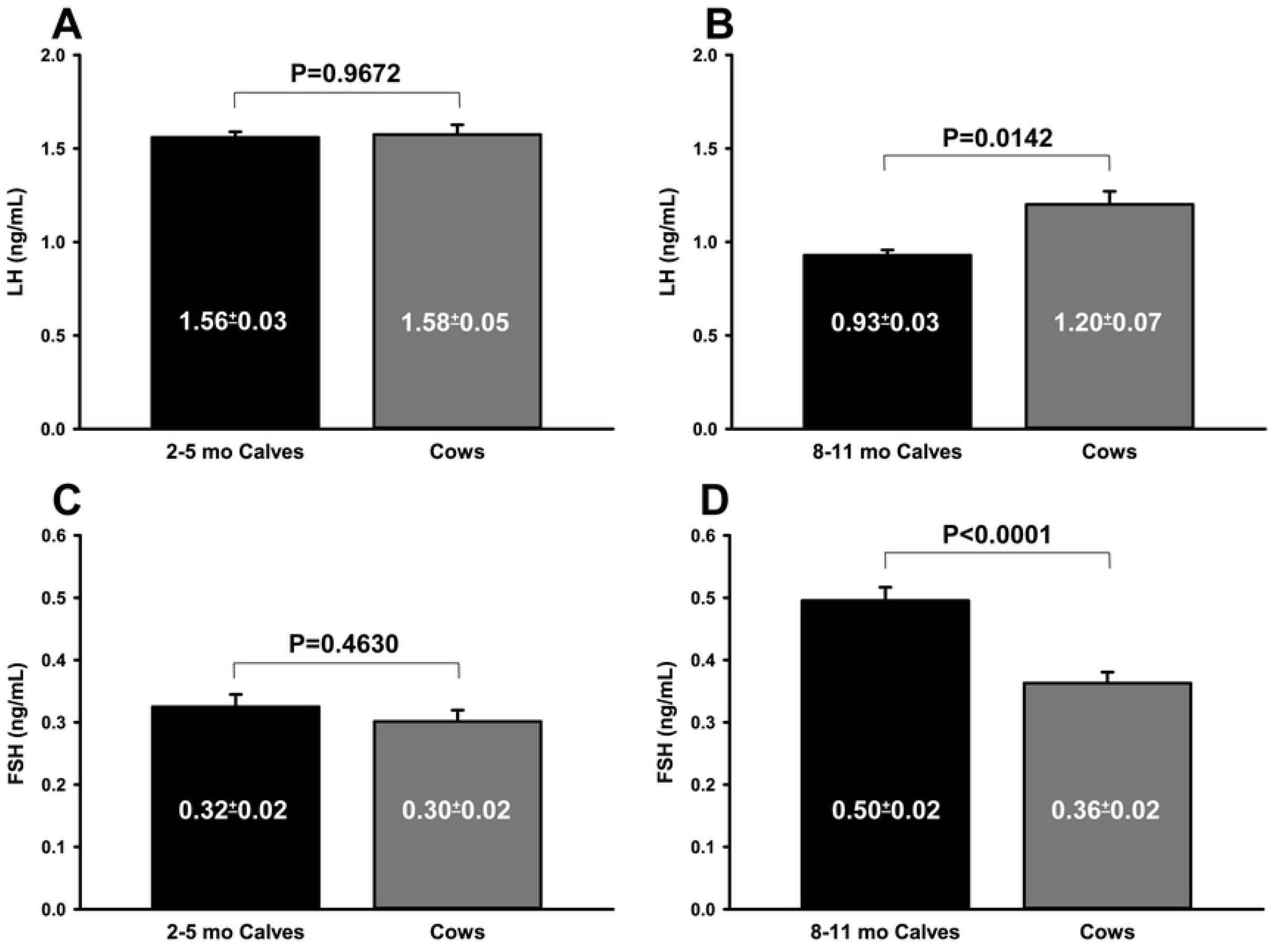
(A-D). Comparison of plasma LH (A, B) and FSH (C, D) concentrations (ng/mL) in Nelore calves during two prepubertal periods (black bars) and non-pregnant, non-lactating pluriparous Nelore cows (gray bars). A: LH, 2- to 5-mo calves vs. cows; B: LH, 8- to 11-mo calves vs. cows; C: FSH, 2- to 5-mo calves vs. cows; D: FSH, 8- to 11- mo calves vs. cows. P values were determined using T-test (A, B) or Mann-Whitney test (C, D). Results are shown as mean ± SEM.

The correlation matrix for reproductive and biometric endpoints is shown in Fig 2 (A, B). During the EPP, moderate to high positive correlations were observed among ovarian diameter, number of follicles > 4.0 mm, diameter of the largest follicle, uterine horn diameter, and rump biometric endpoints (R values ranging from 0.30 to 0.77, P < 0.05). On the other hand, during IPP, most of these associations were weak and non-significant (P > 0.05). In both periods, AFC and the number of follicles ≤ 4 mm only presented a significant positive correlation with ovarian diameter, whereas in the IPP AFC, the number of follicles ≤ 4 mm was negatively correlated with rump width at the ischium, rump area, and body weight (R = −0.10, −0.16, and −0.10, respectively, P < 0.05).

**Fig 2.**
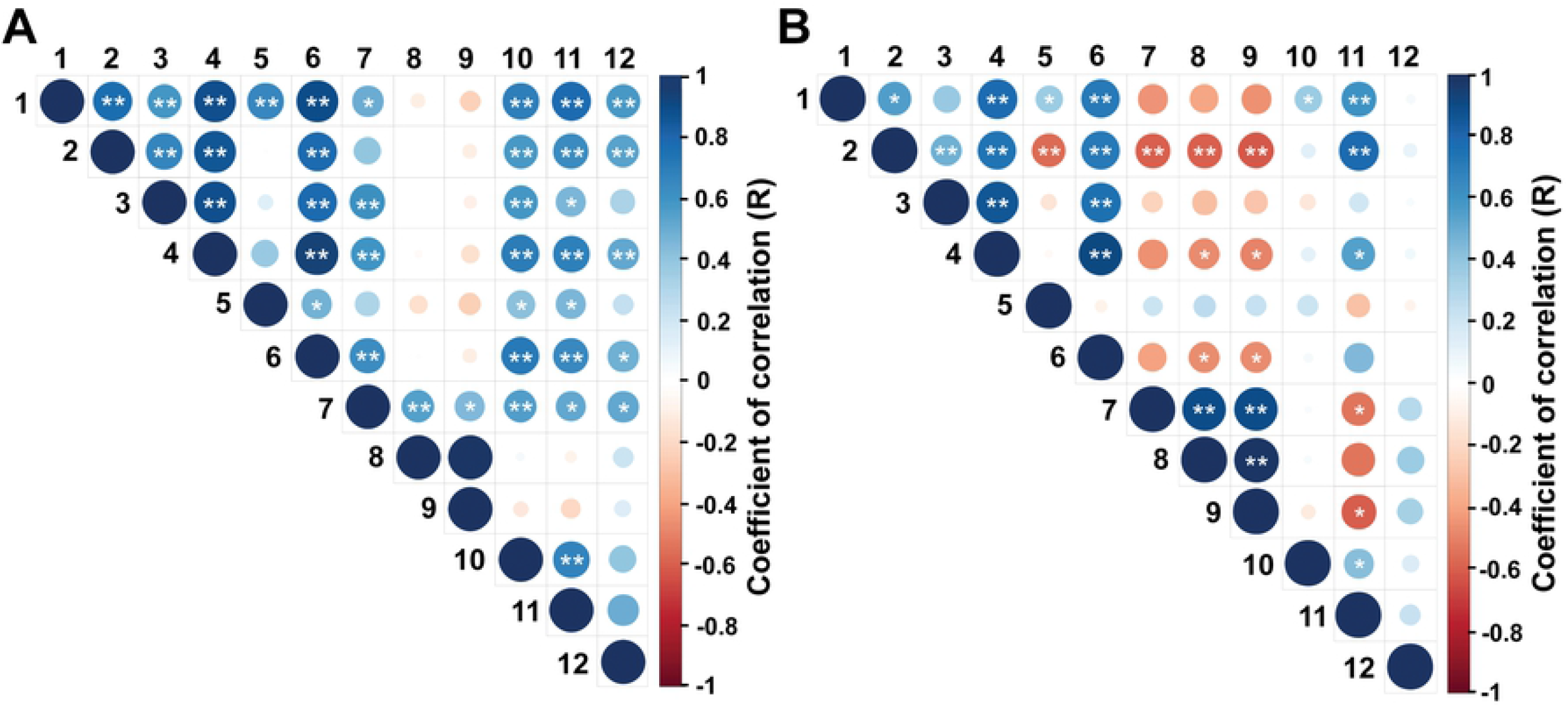
(A-B). Spearman’s correlation between reproductive and biometric parameters in Nelore calves at A: 2 to 5 mo and B: 8 to 11 mo. Endpoints evaluated: 1: rump width at the ilium (cm), 2: rump width at the ischium (cm), 3: rump length (cm), 4: rump area (cm^2^), 5: ratio between ilium’s and ischium’s length, 6: body weight (Kg), 7: ovarian diameter (mm), 8: antral follicle count, 9: number of follicles ≤ 4.0 mm, 10: number of follicles > 4.0 mm, 11: diameter of the largest follicle (mm), 12: uterine horn diameter (mm). * is P < 0.05 and ** is P < 0.01.

The Gompertz model was used to analyze whether the differences observed between correlations in the EPP and IPP were caused by differences in the rate of reproductive development. Despite of the relatively weak values for r^2^ (ranging from 0.2 to 0.4), we could determine the reproductive developmental rate (“a” value in Gompertz equation) for calves at this early age (Fig 3A). During this period, the ovarian diameter and the number of follicles > 4.0 mm increased by 33%, the diameter of the largest follicle increased by 24%, and the uterine horn diameter increased by 19%. To ensure that the slope of the curves was positive, they were compared to a straight line. The curves for all endpoints, except follicles ≤ 4.0 mm, differed (P < 0.05) from the straight line, i.e., represented growing trends (Table 2). Conversely, during the IPP, the equations for the same endpoints showed r^2^ values below 0.01 (Fig 3B), and the curves did not differ from straight lines (P> 0.05, Table 2). The following equations were calculated for the EPP:

Ovarian diameter: 23.57*exp(−1.00983*exp(−0.33040*x)); r^2^ = 0.44
Follicles > 4.0 mm: 8*exp(−2.52892 *exp(−0.33549*x)); r^2^ = 0.23
Diameter of the largest follicle: 11.30*exp(−0.99893*exp(−0.24732*x)); r^2^ = 0.21
Uterine horn diameter: 11.20*exp(−0.53403*exp(−0.19523*x)); r^2^ = 0.16
Rump area: 711.40 * exp (−2.05001 * exp (−0.64280 * x)); r^2^ = 0.748

The following equations were calculated for the IPP:

Ovarian diameter: 23.65*exp(−0.43306*exp(−0.05201*x)); r^2^ = 0.009
Follicles > 4.0 mm: 14*exp(−0.84750*exp(−0.02280*x)); r^2^ = 0.002
Diameter of the largest follicle: 10.65*exp(−0.62063*exp(−0.08197*x)); r^2^ = 0.014
Uterine horn diameter: 13.15*exp(−0.55009*exp(−0.08051*x)); r^2^ = 0.024
Rump area: 1026.22 * exp (−3.7862 * exp (−0.3416 * x)); r^2^ = 0.303

**Fig 3.**
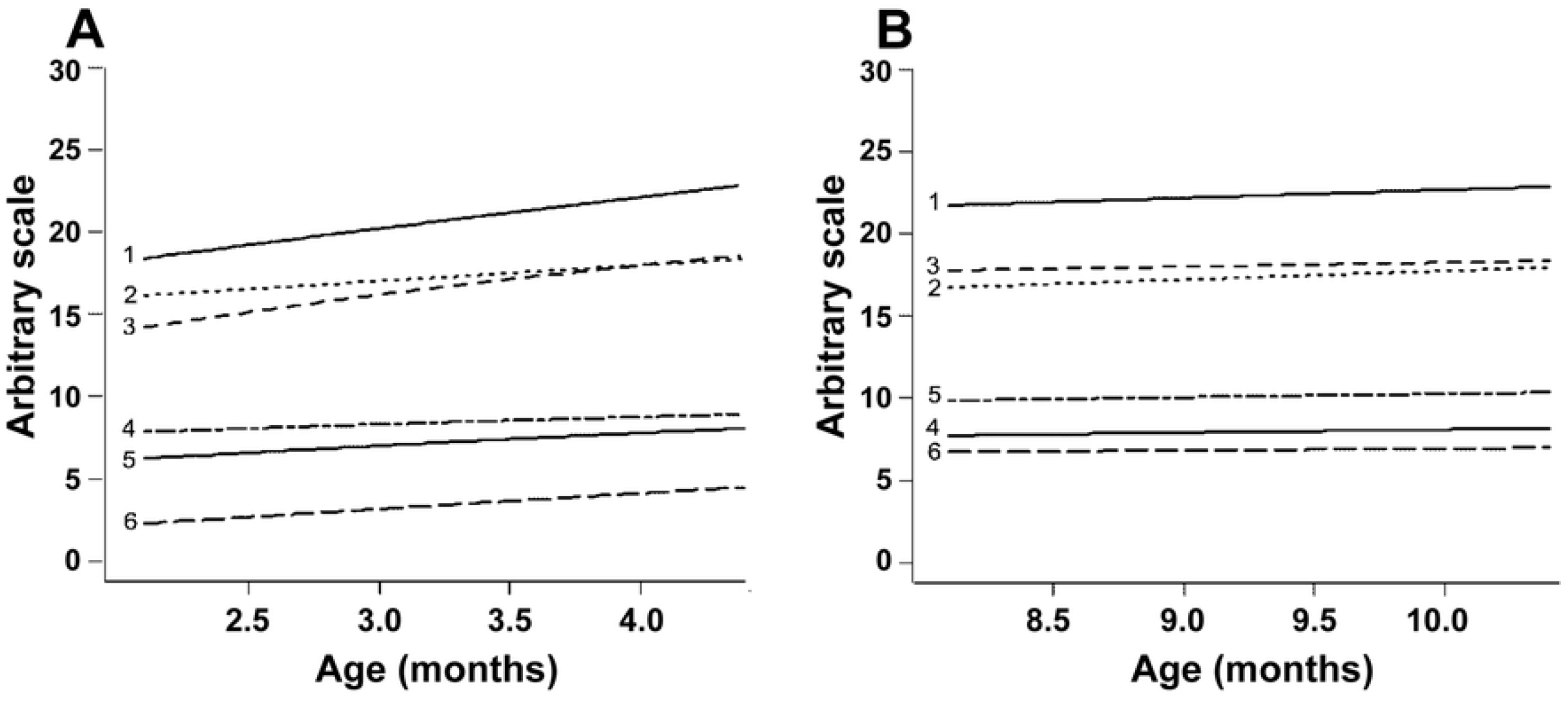
(A-B). Gompertz curves for reproductive parameters in Nelore calves from 2 to 5 mo (A) and from 8 to 11 mo (B). The parameters evaluated and respective r^2^ values at the periods A and B, respectively, are as follows: 1: antral follicle count (r^2^ = 0.056 and 0.001), 2: follicles ≤ 4.0 mm (r^2^ = 0.014 and 0.002), 3: ovarian diameter (r^2^ = 0.442 and 0.009), 4: diameter of the largest follicle (ŕ = 0.207 and 0.014), 5: uterine horn diameter (r^2^ = 0.157 and 0.024), and 6: ovarian follicles > 4.0 mm (r^2^ = 0.232 and 0.001). The R software output presents mean values in the Y axis in an arbitrary scale so that all endpoints fit in the same picture.

**Table 2.** Comparison between the Gompertz curves for reproductive parameters (referred to in Fig 3) and a straight line in 2- to 5- and 8- to 11-month-old Nelore calves. Results are shown as sum of squares (SQ), F, and P-values. When P < 0.05, the curve slope was considered positive.

| Endpoint | Age group |  |  |  |  |  |
| --- | --- | --- | --- | --- | --- | --- |
|  | 2 to 5 months |  |  | 8 to 11 months |  |  |
|  | SQ | F | P-value | SQ | F | P-value |
| Ovary diameter (mm) | -238.9 | 74.6 | <0.001 | -4.3 | 0.9 | 0.3 |
| AFC (n) | -244.0 | 5.6 | 0.02 | -16.3 | 0.1 | 0.6 |
| Follicles $\leq 4.0$ mm (n) | -61.1 | 1.3 | 0.26 | -19.8 | 0.2 | 0.6 |
| Follicles $> 4.0$ mm (n) | -60.1 | 28.4 | <0.001 | -0.9 | 0.2 | 0.6 |
| Largest follicle (mm) | -42.3 | 24.5 | <0.001 | -2.4 | 1.5 | 0.2 |
| Uterine diameter (mm) | -15.2 | 17.5 | <0.001 | -2.9 | 2.6 | 0.1 |

Changes in rump geometry are shown in Fig 4 (A-D). The non-linear regression analysis demonstrated a greater developmental rate in rump area during EPP, which increased by 64% (r^2^ = 0.75) compared to 34% (r^2^ = 0.31) during the IPP (Fig 5 A, B). During the EPP, there was also a greater change in rump geometry, which changed from a more squared to a trapezoidal shape (Fig 4 A, B). The average rump area was 1,684.2 ± 173,8 cm^2^ for cyclic cows (control) and 514.2 ± 101.4 cm^2^ and 856.72 ± 79.1 cm^2^ for EPP and IPP calves, respectively. The ratio rump width at ilium: at ischium differed between EPP and IPP (P < 0.05). The calves reached puberty at an average of 20.5 mo (ranging from 17 to 24 mo).

**Figure 4.**
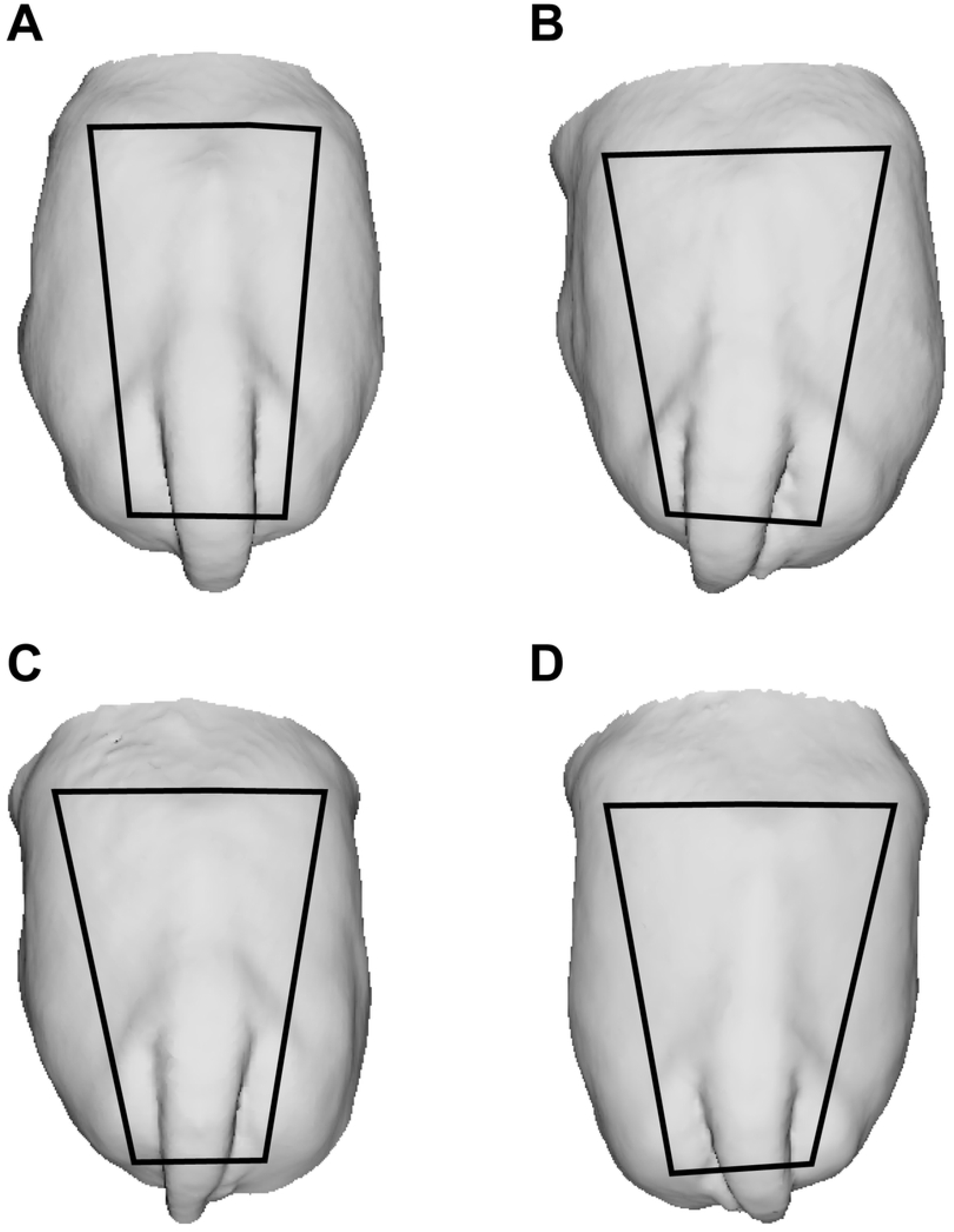
(A-D). Changes in rump geometry during prepubertal development in Nelore (*Bos indicus*) calves. Back-view 3D images of Nelore females’ rump at 2 (A) and 5 (B) and at 8 (C) and 11 (D) mo. Trapezoids show changes in rump geometry. To build the trapeze, the iliac crest and tuber ischiadicum were used as reference points.

**Figure 5.**
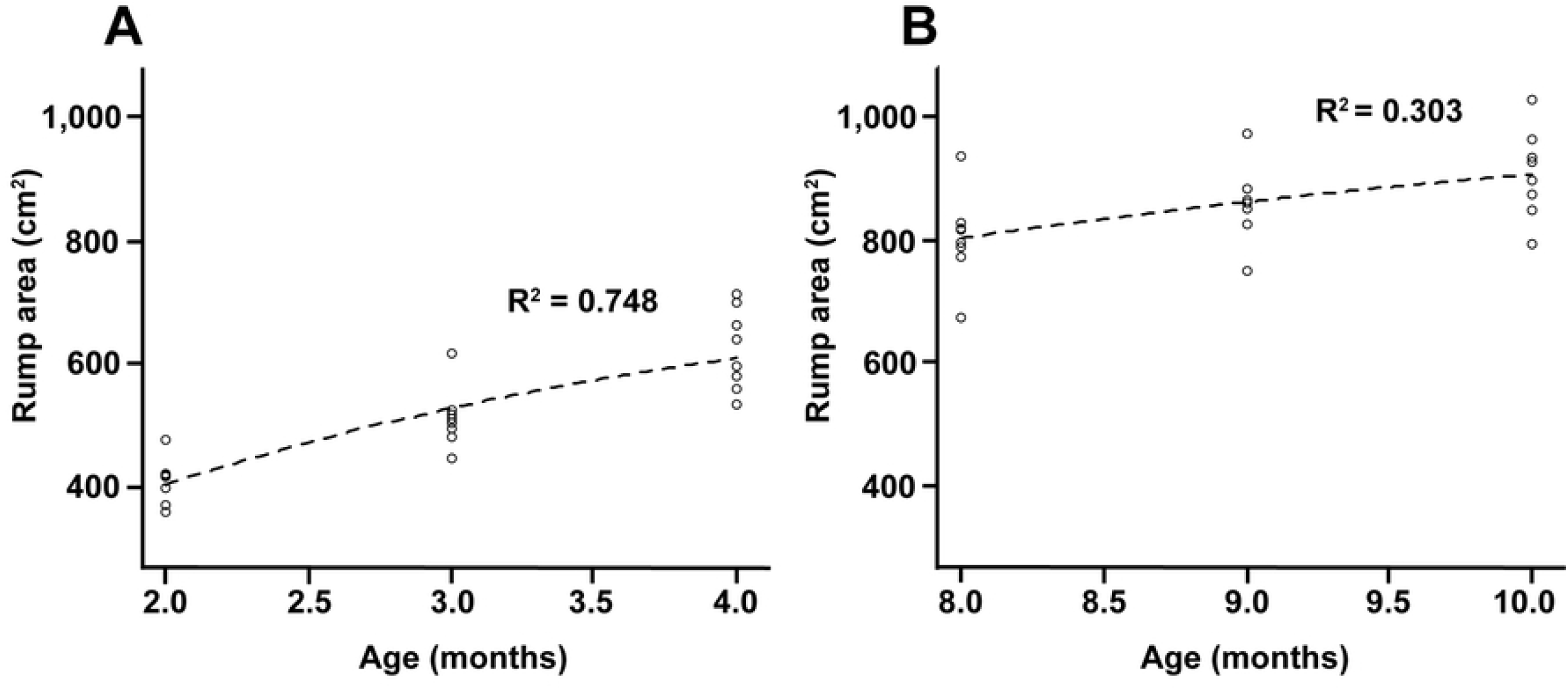
(A-B). Non-linear regression curves with r^2^ values of rump geometry measures in calves at 2 to 5 mo (A) and 8 to 11 mo (B).

## Discussion

The goal of this study was to characterize the reproductive, biometric, and hormonal patterns in the initial and intermediate prepubertal periods in Nelore (*B. indicus*) calves. The present results support the hypothesis that the dynamics of reproductive and biometric development during the prepubertal period differs over time.

There was a progressive increase in ovarian activity in calves, indicated by the difference in the number of growing follicles and by the maximum size of the largest (presumptive dominant) follicle and indirectly by uterine horn diameter between EPP (2 to 5 mo) and IPP (8 to 11 mo) calves. However, the dynamics of ovarian and uterine development differed within periods, with a higher growth rate during the EPP, as demonstrated by the differences in the non-linear Gompertz curves. In EPP calves, there was a significant increase in all endpoints other than AFC, whereas in IPP calves, the curves were not significantly different from a straight line, i.e., there was a relative stabilization in reproductive development. The faster development observed in younger calves is in line with the so-called “mini puberty” previously described in *B. taurus* [1] as well as in sheep [4] and humans [reviewed by 3]. Coherently, ovarian activity showed positive and significant correlations with ovarian and uterine horn diameters as well as with biometric endpoints (length at ilium, length in ischium, rump length, rump area, ratio between rump width at ilium: at ischium, and body weight) during the EPP. On the other hand, most of these correlations became nonexistent or even negative in IPP calves.

The marked changes observed in EPP calves are likely driven by the increasing production of estradiol (not evaluated in this study) by growing follicles. The circulating estradiol promotes both uterine growth [23] and somatic development [5], and ovarian activity at this early age seems to be important for the acquisition of primary and secondary sexual characteristics. In IPP calves, on the other hand, increases in body weight or size were associated with a stabilization rather than proportional changes in ovarian activity. Previous studies have described that, in the first months of life, there is a faster development of the reproductive tract, followed by a plateau and by a resumption of progress near puberty [1, 24]. However, in the current study, this plateau in reproductive development occurred in the period referred to as the intermediate prepubertal period (IPP, 8 to 11 mo). To make a parallel, it would correspond, in precocious *B. taurus* breeds, to the peripubertal period, in which remarkable neuroendocrine and physiological changes are in progress, with concurrent changes in the reproductive tract. The age at puberty in Holstein calves is, on average, 250 days (8.3 mo), with approximately 30% of the calves reaching puberty as early as at 7 mo [25]. The Angus, a beef breed, reaches puberty at about 13 mo [26]. Conversely, our calves attained puberty at an average of 20.5 mo, consistently with the age previously reported by others for the Nelore breed [18]. In this regard, the present study demonstrates the differences in the timing of the events in prepubertal *B. indicus* calves, with the plateau phase extending to the period here defined as the IPP.

It is noteworthy that changes in rump geometry (from squared to trapezoid) were observed only in EPP calves, and further development was characterized mainly by increases in rump size rather than in shape. Rump characteristics are associated with pregnancy failure, anestrus, and dystocia [27–29] in heifers, highlighting the importance of the changes in rump development that take place during the early prepubertal phase.

A novelty of the current study was the use of 3D image technologies to demonstrate these changes in rump geometry. Structured light scanning allows the fast and accurate generation of 3D models from cattle, which can be freely rotated in a 3D virtual space and used to define reference marks and acquire biometric measurements [30]. In the current study, rump size and geometry were calculated using as references the prominences of the *tuber ischiae* and the *tuber coxae* of the pelvic bones, which can be easily spotted in the 3D models.

Differently from the other ovarian endpoints evaluated, the number of sonographic-detectable antral follicles (AFC) did not differ between age groups, and analysis by the Gompertz curve showed little or no trend of increase in AFC values within the periods analyzed. Coherently, in both periods, AFC was positively correlated only with ovarian diameter and, in older calves, negatively correlated with rump development and body weight. The association of AFC and ovary size has previously been described in pubertal cattle [31], and in the current study, we now show that this relationship also occurs in early prepubertal calves. As AFC presents a high individual repeatability from prepubertal to post-pubertal heifers [32], our findings suggest that oocyte donor selection can be performed in young calves, with no expected effect on other characteristics associated with prepubertal sexual development. However, the association between AFC and somatic development remains to be elucidated.

The progressive increase in the number of growing follicles and in the maximum size of the largest follicle observed in this study in the EPP calves was consistent with the hypothesis that a higher hypophyseal release of FSH and LH occurs during early calfhood. In fact, plasma concentrations of FSH and LH in EPP calves were similar to those in control cows. The hypophyseal gonadotropins support follicle growth and steroidogenesis (reviewed by [33]), which in turn may have promoted uterine development and the changes in rump geometry, as observed in the current study. Mauras et al. [34] observed that, in children, the increase in GnRH pulse amplitude leads to a greater FSH and LH release and subsequent estradiol production, which in turn increases the production of growth hormone (GH) and insulin-like growth hormone (IGF-1), as well as calcium absorption and skeleton mineralization.

On the other hand, the events observed during IPP support the hypothesis of an increased sensitivity of the hypothalamic-hypophyseal axis as a response to the negative feedback from estradiol. Previous studies have shown that after the transient hypothalamic-pituitary-gonadal axis activation soon after birth, there is a decrease in plasma concentration of gonadotropins [1, 4]. In fact, in our study, LH concentrations were lower in IPP calves than in control cows. The lower amplitude and frequency of LH pulses may have reduced estradiol production and, thus, the rate of development of primary and secondary sexual characteristics. Moreover, the lower negative feedback on FSH production [35] would explain the plasma concentrations of FSH in IPP calves, which were higher than those in the control cows.

## Conclusions

The EPP in Nelore calves is characterized by increased ovarian activity, uterine development, and rump geometry change, contrasting with the relative stabilization observed in the IPP. Additionally, the plateau phase in Nelore calves extends to a moment when *B. taurus* breed calves are already in the peripubertal period. These findings highlight the importance of the early prepubertal period for the development of primary and secondary characteristics and shed light on the implications of the differences in physiology between these periods for the use of calves as oocyte donors.

## Acknowledgements

The authors thank the Federal University of Uberlandia - Veterinary Sciences Graduate Program (PPGCV-UFU) and the Dean of Research and Graduation (PROPP), the Brazilian Agricultural Research Corporation (Embrapa), and the collaboration of Embrapa Sucupira Experimental Farm personnel, Brasilia-DF, Brazil. We also thank Dr. Guilherme de Paula Nogueira’s team at the Laboratory of Endocrinology (Sao Paulo State University - UNESP, Aracatuba, SP, Brazil) for the LH and FSH analyses, and Joseane Padilha da Silva (Embrapa Genetic Resources and Biotechnology) for the statistical analyses

## Financial support

This research was supported by FAP-DF (Federal District Research Support Foundation. Projects # 193.001.393/2016 and 193.001.640/2017) and it was partially supported by the Coordination for the Improvement of Higher Education Personnel - Brazil (CAPES) - Finance Code 001.

## Notes

### Competing Interest Statement

The authors have declared no competing interest.

## References

[1] Rawlings NC, Evans ACO, Honaramooz A, Bartlewski PM. Antral follicle growth and endocrine changes in prepubertal cattle, sheep and goats. Anim Reprod Sci. 2003; 78: 259–270. DOI: 10.1016/s0378-4320(03)00094-0

[2] Hernandez-Medrano JH, Campbell BK, Webb R. Nutritional influences on folliculogenesis. Reprod Domest Anim. 2012; 47: 274–282. DOI: 10.1111/j.1439-0531.2012.02086.x

[3] Kuiri-Hänninen T, Sankilampi U, Dunkel L. Activation of the hypothalamic-pituitary-gonadal axis in infancy: minipuberty. Horm Res Paediatr. 2014; 82: 73–80. DOI: 10.1159/000362414

[4] Torres-Rovira L, Succu S, Pasciu V, Manca ME, Gonzalez-Bulnes A, Leoni GG, et al. Postnatal pituitary and follicular activation: a revisited hypothesis in a sheep model. Reproduction. 2016; 151: 215–225. DOI: 10.1530/REP-15-0316

[5] Connelly KJ, Larson EA, Marks DL, Klein RF. Neonatal estrogen exposure results in biphasic age-dependent effects on the skeletal development of male mice. Endocrinology. 2015; 156: 193–202. DOI: 10.1210/en.2014-1324

[6] Smith MF, Geisert RD, Parrish JJ. Reproduction in domestic ruminants during the past 50 yr: discovery to application. J Anim Sci. 2018; 96: 2952–2970. DOI: 10.1093/jas/sky139

[7] Oursler MJ, Cortese C, Keeting P, Anderson MA, Bonde SK, Riggs BL, Spelsberg TC. Modulation of transforming growth factor-β production in normal human osteoblast-like cells by 17 β-estradiol and parathyroid hormone. Endocrinology. 1991; 129: 3313–3320. DOI: 10.1210/endo-129-6-3313

[8] Michael H, Härkönen PL, Väänänen HK, Hentunen TA. Estrogen and testosterone use different cellular pathways to inhibit osteoclastogenesis and bone resorption. J Bone Miner Res. 2005; 20: 2224–2232. DOI: 10.1359/JBMR.050803

[9] Day ML, Anderson LH. Current concepts on the control of puberty in cattle. J Anim Sci. 1998; 76:1–15. DOI: 10.2527/1998.76suppl_31x

[10] Presicce GA, Jiang S, Simkin M, Zhang L, Looney CR, Godke RA, et al. Age and hormonal dependence of acquisition of oocyte competence for embryogenesis in prepubertal calves. Biol Reprod. 1997; 56: 386–392. DOI: 10.1095/biolreprod56.2.386

[11] Brogliatti GM, Adams GP. Ultrasound-guided transvaginal oocyte collection in prepubertal calves. Theriogenology. 1996; 45: 1163–1176. DOI: 10.1016/0093-691x(96)00072-6

[12] Majerus V, De Roover R, Etienne D, Kaidi S, Massip A, Dessy F, et al. Embryo production by ovum pick up in unstimulated calves before and after puberty. Theriogenology. 1999; 52: 1169–1179. DOI: 10.1016/S0093-691X(99)00209-5

[13] Jiang J, Ma L, Prakapenka D, VanRaden PM, Cole JB, Da Y. A large-scale genome-wide association study in US Holstein cattle. Front genet. 2019; 10: 412. DOI: 10.3389/fgene.2019.00412

[14] Baldassarre H. Laparoscopic ovum pick-up followed by *In vitro* embryo production and transfer in assisted breeding programs for ruminants. Animals. 2021; 11: 216. DOI: 10.3390/ani11010216

[15] Batista EOS, Guerreiro BM, Freitas BG, Silva JCB, Vieira LM, Ferreira RM, et al. Plasma anti-Müllerian hormone as a predictive endocrine marker to select *Bos taurus* (Holstein) and *Bos indicus* (Nelore) calves for *in vitro* embryo production. Domest Anim Endocrinol. 2016; 54: 1–9. DOI: 10.1016/j.domaniend.2015.08.001

[16] Baldassarre H, Currin L, Michalovic L, Bellefleur AM, Gutierrez K, Mondadori RG, et al. Interval of gonadotropin administration for in vitro embryo production from oocytes collected from Holstein calves between 2 and 6 months of age by repeated laparoscopy. Theriogenology. 2018; 116: 64–70. DOI: 10.1016/j.theriogenology.2018.05.005

[17] Bernal-Ulloa SM, Heinzmann J, Herrmann D, Hadeler KG, Aldag P, Winkler S, et al. Cyclic AMP affects oocyte maturation and embryo development in prepubertal and adult cattle. Plos One. 2016; 11(2): e0150264. DOI: 10.1371/journal.pone.0150264

[18] Monteiro FM, Mercadante MEZ, Barros CM, Satrapa RA, Silva JAV, Oliveira LZ, et al. Reproductive tract development and puberty in two lines of Nellore heifers selected for postweaning weight. Theriogenology. 2013; 80: 10–17. DOI: 10.1016/j.theriogenology.2013.02.013

[19] Abeni F, Petrera F, Le Cozler Y. Effects of feeding treatment on growth rates, metabolic profiles and age at puberty, and their relationships in dairy heifers. Animal. 2019; 13: 1020–1029. DOI: 10.1017/S1751731118002422

[20] Sartori R, Gimenes LU, Monteiro Jr PL, Melo LF, Baruselli OS, Bastos MR. Metabolic and endocrine differences between *Bos taurus* and *Bos indicus* females that impact the interaction of nutrition with reproduction. Theriogenology. 2016; 86: 32–40. DOI: 10.1016/j.theriogenology.2016.04.016

[21] Nogueira GP. Puberty in South American Bos indicus (Zebu) cattle. Anim Reprod Sci. 2004; 82-83: 361–372. DOI: 10.1016/j.anireprosci.2004.04.007

[22] Bolt DJ, Scott V, Kiracofe GH. Plasma LH and FSH after estradiol, norgestomet and GnRH treatment in ovariectomized beef heifers. Anim Reprod Sci. 1990; 23: 263–271. DOI: 10.1016/0378-4320(90)90040-M

[23] Jdidi H, Kouba FG, Aoiadni N, Abdennabi R, Turki M, Makni-Ayadi F, et al. Effects of estrogen deficiency on liver function and uterine development: assessments of *Medicago sativa’s* activities as estrogenic, anti-lipidemic, and antioxidant agents using an ovariectomized mouse model. Arch Physiol Biochem. 2019; 18: 1–12. DOI: 10.1080/13813455.2019.1625927

[24] Honaramooz A, Aravindakshan J, Chandolia RK, Beard AP, Bartlewski PM, Pierson RA, et al. Ultrasonographic evaluation of the pre-pubertal development of the reproductive tract in beef heifers. Anim Reprod Scie. 2004; 80: 15–29. DOI: 10.1016/S0378-4320(03)00136-2

[25] Bruinjé TC, Rosadiuk JP, Moslemipur F, Sauerwein H, Steele MA, Ambrose DJ. Differing planes of pre- and postweaning phase nutrition in Holstein heifers: II. Effects on circulating leptin, luteinizing hormone, and age at puberty. J Dairy Sci. 2021; 104: 1153–1163. DOI: 10.3168/jds.2020-18810

[26] Allen CC, Tedeschi LO, Keisler DH, Cardoso RC, Alves BRC, Amstalden M, et al. Interaction of dietary energy source and body weight gain during the juvenile period on metabolic endocrine status and age at puberty in beef heifers. J Anim Sci. 2017; 95: 2080–2088. DOI: 10.2527/jas.2016.1002

[27] Wall E, White IMS, Coffey MP, Brotherstone S. The relationship between fertility, rump angle, and selected type information in Holstein-Friesian cows. J Dairy Sci. 2005; 88: 1521–1528. DOI: 10.3168/jds.S0022-0302(05)72821-6

[28] Holm DE, Webb EC, Thompson PN. A new application of pelvis area data as culling tool to aid in the management of dystocia in heifers. J Anim Sci. 2014; 92: 2296–2303. DOI: 10.2527/jas.2013-6967

[29] Holm DE, Nielen M, Jorritsma R, Irons PC, Thompson PN. Ultrasonographic reproductive tract measures and pelvis measures as predictors of pregnancy failure and anestrus in restricted bred beef heifers. Theriogenology. 2016; 85: 495–501. DOI: 10.1016/j.theriogenology.2015.09.031

[30] Viana JHM, Hinduja S, Bártolo PJS. Estimation of biometric parameters from cattle rump using free-hand scanning and a 3D data processing algorithm. Virtual Phys Prototyp. 2016; 11: 167–172. DOI: 10.1080/17452759.2016.1211292

[31] Ireland JJ, Smith GW, Scheetz D, Jimenez-Krassel F, Folger JK, Ireland JL, et al. Does size matter in females? An overview of the impact of the high variation in the ovarian reserve on ovarian function and fertility, utility of anti-Müllerian hormone as a diagnostic marker for fertility and causes of variation in the ovarian reserve in cattle. Reprod Fert Develop. 2011; 23: 1–14. DOI: 10.1071/RD10226

[32] Morotti F, Santos GMG, Júnior CK, Silva-Santos KC, Roso VM, Seneda MM. Correlation between phenotype, genotype and antral follicle population in beef heifers. Theriogenology. 2017; 91: 21–26. DOI: 10.1016/j.theriogenology.2016.12.025

[33] Kinder JE, Bergfeld EG, Wehrman ME, Peters KE, Kojima FN. Endocrine basis for puberty in heifers and ewes. J Reprod Fert. 1995; 49: 393–407. PMID: 7623330

[34] Mauras N, Rogol AD, Haymond MW, Veldhuis JD. Sex steroids, growth hormone, insulin-like growth factor-1: neuroendocrine and metabolic regulation in puberty. Horm Res Paediatr. 1996; 45: 74–80. DOI: 10.1159/000184763

[35] Ginther OJ, Bergfelt DR, Kulick LJ, Kot K. Selection of the dominant follicle in cattle: establishment of follicle deviation in less than 8 hours through depression of FSH concentrations. Theriogenology. 1999; 52:1079–1093. DOI: 10.1016/S0093-691X(99)00196-X

